# Genetic and environmental influences on the distributions of three chromosomal drive haplotypes in maize

**DOI:** 10.1101/2025.05.22.655462

**Authors:** Meghan J. Brady, R. Kelly Dawe

## Abstract

Meiotic drive elements are regions of the genome that are transmitted to progeny at frequencies that exceed Mendelian expectations, often to the detriment of the organism. In maize there are three prevalent chromosomal drive elements known as Abnormal chromosome 10 (Ab10), K10L2, and the B chromosome. There has been much speculation about how these drivers might interact with each other and the environment in traditional maize landraces and their teosinte ancestors. Here we used genotype-by-sequencing data to score more than 10,000 maize and teosinte lines for the presence or absence of each driver. Less than ∼0.5% of modern inbred lines carry chromosomal drivers. Among individuals from 5331 open-pollinated landraces, 6.32% carried Ab10, 5.16% carried K10L2, and 12.28% carried at least one B chromosome. Using a GWAS approach we identified unlinked loci that associate with the presence or absence of the selfish genetic elements. Many genetic modifiers are positively associated with the drivers, suggesting that there may have been selection for alleles that ameliorate their negative fitness consequences. We then assessed the contributions of population structure, associated loci, and the environment on the distribution of each chromosomal driver. There was no significant relationship between any chromosomal driver and altitude, contrary to conclusions based on smaller studies. Our data suggest that the distribution of the major chromosomal drivers is primarily influenced by neutral processes and the deleterious fitness consequences of the drivers themselves. While each driver has a unique relationship to genetic background and the environment, they are largely unconstrained by either.

**AUTHOR SUMMARY:** Meiotic drivers are selfish genetic elements that are transmitted to progeny at frequencies that exceed what is expected by chance. There are three such meiotic drivers in maize known as Ab10, K10L2 and the B chromosome. Prior data indicate that each driver lowers plant fitness to some degree, partially explaining the fact that they are usually observed at low frequencies, but the impact of environment and genetic modifiers has not been investigated. Here we identified the presence or absence of each driver using low coverage sequencing data from over 10,000 maize inbreds, open pollinated landraces, and accessions of teosinte -- the grass-like ancestor of maize. We found that they exist at 5-12% frequencies in open pollinated varieties, but are rare in modern inbreds. We went on to use location data to assess the impact of environmental variables on their distribution, and applied genome-wide association methods to look for genetic modifiers that might contribute to their prevalence. We found that neither the environment nor genetic modifiers have strong influences on the distribution of the three meiotic drive elements, suggesting that their low but consistent frequencies are determined primarily by the fitness consequences of the drivers themselves.

## INTRODUCTION

While most genes in most species are transmitted in predictable Mendelian patterns, there are striking exceptions. Genes, gene complexes, sections of chromosomes, and entire chromosomes have evolved mechanisms that ensure they are transmitted at higher frequencies than would be expected based on chance [1–4]. These genetic elements generally confer no selective advantages to the species, and are often deleterious, but are nevertheless maintained in populations based on their selfish properties. They are often described with the catch-all term meiotic drive [5], though only a subset of meiotic drivers affect meiosis. Meiotic drivers that manipulate meiosis often do so in species where female meiosis results in only one functional egg cell. An example is the preferential transmission of larger centromeres towards the egg cell in some mice lines [6]. A more common class of meiotic driver interferes with the function of male gametes, often by setting up a dynamic where sperm or pollen are killed by a toxin unless they inherit the antidote present on the driving chromosome [7]. The term meiotic drive is also used to describe the maintenance of supernumerary B chromosomes [8], as well as many other varied phenomena, including biased gene conversion processes [9], mobile toxin-antidote systems [10], and engineered gene drive systems based on CRISPR [11].

The maize genome contains at least three meiotic drive elements that distort transmission of large chromosomal regions: Abnormal Chromosome 10 (Ab10), K10L2, and the B chromosome. Ab10 is a large variant of normal chromosome 10 (N10) that acts as a female meiotic driver. Approximately 14% of the maize genome is composed of tandem repeats arrays called knobs [12]. They come in two classes defined by their repeat element: TR-1 and knob180. The Ab10 haplotype contains knobs of both classes with the knob180 knob being one of the largest knobs in the genome [13,14]. The Ab10 haplotype encodes two kinesin proteins, KINDR and TRKIN, which interact with knob180 and TR-1 knobs respectively [15,16]. The kinesins pull the knobs ahead of the centromeres during meiotic anaphase resulting in their preferential transmission to the egg cell [17,18]. By this mechanism the Ab10 haplotype as well as knobs throughout the genome are preferentially transmitted to ∼60-80% of the next generation [18]. Ab10 is present in ∼5% of maize landraces [19,20], but is prevented from going to fixation because it impairs fitness when homozygous [17,21]. Ab10 is recognized as an important driver of maize genome evolution [15,22].

The K10L2 variant is similar to Ab10 but is much smaller, having only two TR-1 rich knobs and the *Trkin* gene. It shows 51-52% meiotic drive when paired with normal chromosome 10 (N10) [23]. When K10L2 is paired with Ab10, Ab10 drive is severely suppressed, demonstrating K10L2 not only can drive itself but compete with the stronger Ab10 drive system. Recent data demonstrate that the region between the two TR-1-rich knobs is similar in sequence to a portion of at least one structural variant of Ab10 (Ab10 Type I), suggesting that Ab10 may have subsumed the K10L2 haplotype in recent evolutionary history [14]. K10L2 is present in about 5% of maize landraces [19,20], but the fact that at least one traditional inbred line is homozygous for K10L2 suggests that it may not have severe effects on fitness when homozygous [23].

The B chromosome is a ∼150 Mb supernumerary chromosome composed primarily of transposable elements (TEs), organellar sequences, and a B-specific repeat element [24]. The B chromosome can accumulate to high copy numbers by a mechanism that takes advantage of the fact that there are two sperm in each pollen grain; one fertilizes the egg cell and the other fertilizes the central cell that gives rise to the starchy endosperm. The B chromosome normally non-disjoins at the second pollen mitosis and the sperm carrying two copies of the B chromosome preferentially fertilizes the egg [25]. There is known variation among lines for the efficiency of the second step. Most lines allow sperm carrying the B chromosome to preferentially fertilize the egg, but multiple lines do not [8], or even reverse it, such that B chromosome-carrying sperm preferentially fertilize the central cell [26]. In natural populations, B chromosomes are found in about 10% of landraces with copy numbers that are usually between 1-3 but may be as high as 14 [19,20,27]. When B chromosomes are present at higher than 15 copies the plants display reduced seed set and pollen viability [28].

The factors affecting the frequency and distribution of maize chromosomal drivers are not well understood. A prior modeling study showed that the known fitness defects associated with Ab10 can largely, but not completely, explain its low frequency in landraces and teosintes [21], suggesting that there are other variables that influence its distribution. Likely influences include the environment and genetic modifiers [2,29]. Prior evidence suggests that altitude may influence the distribution of both Ab10 and the B chromosome [22,27,30–33], which may be related to the fact that high-altitude maize lines tend to have smaller genomes [33–35]. Genetic variation outside of the drive haplotypes is also expected to alter their frequencies. Alleles that reduce the efficiency of drive (suppressors) should be selected for when the fitness consequences of the drive haplotypes are high [2,29,36–38]. It is possible that the resistance of some lines to the preferential fertilization of B chromosomes reflects ongoing selection against high B chromosome copy numbers [8]. The three maize drivers may also interact with each other in unpredictable ways. For instance, in some backgrounds, the B chromosome causes the breakage of chromosomes at knobs, including Ab10 [39,40].

Here we developed methods to detect maize chromosomal drive haplotypes in genotype-by-sequencing (GBS) data, and scored their presence or absence in ∼10,000 maize inbred, maize landrace, and teosinte individuals. We then determined how their distribution relates to genetic background, population structure, and the environment. In open pollinated landraces and teosintes, their distributions are significantly influenced by genetic modifiers and environmental factors, though the effects are small. The combined data, and the fact that all three drivers are present at very low frequencies in modern inbred lines, support the view that the major limitation to their spread is the negative fitness consequences of the drivers themselves.

## RESULTS

### Genotype-by-sequencing data can reliably detect large structural variants

We speculated that the low coverage sequence data from GBS studies might be useful for identifying large chromosomal drive haplotypes (referred to as CDH in this study). To test the feasibility of this idea, we first generated GBS data from a collection of control lines carrying different isolates of Ab10, K10L2, and B chromosomes, as well as associated controls. The GBS data was then mapped to reference genomes carrying each drive haplotype: Ab10 [41], K10L2 [14], and the B chromosome [24]. We then computed a tag index, which is a function of both the number of tag sites and the read depth over tag sites, in 1 Mb windows (Fig S1A).

When the tag index was plotted as a heat map, the presence or absence of a CDH became visually apparent (Fig 1A). We then automated the scoring of CDHs using an iterative k-means clustering approach (Fig 1A,1B) and achieved 100% accurate discrimination of the presence and absence of each CDH in our control dataset. To estimate CDH copy number, we normalized CDH tag depth by the average tag depth across all single copy core genes [12] and correctly identified all Ab10 and K10L2 homozygous samples in our control set (Fig 2C). We applied the same method to estimate the copy numbers of B chromosomes, though in this case we did not know the copy number in our control samples (so we refer to our estimates as pseudo-copy number).

**Figure 1.**
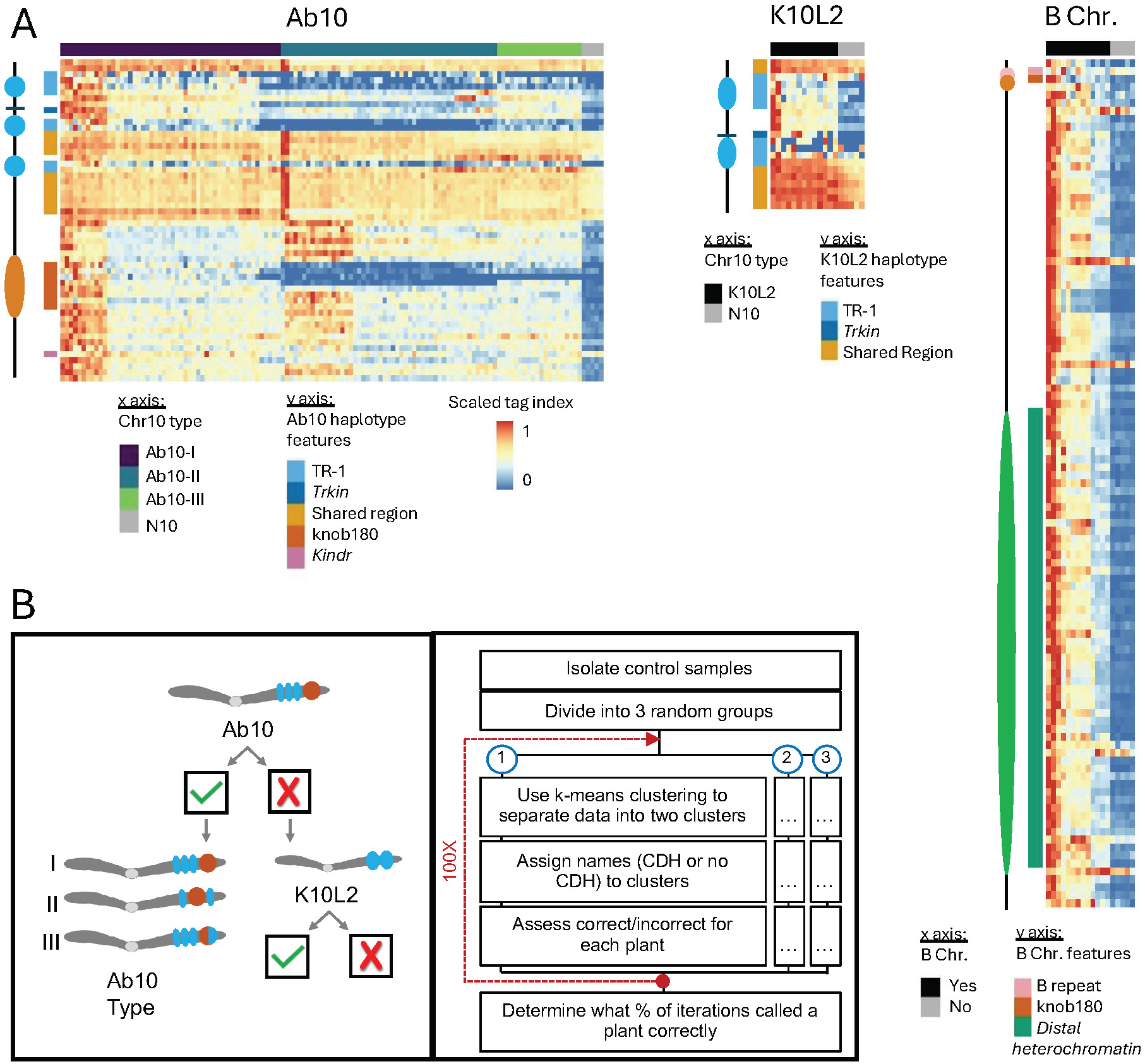
Detection of CDHs in GBS data. **A**. Min/max scaled tag index for Ab10, K10L2, and the B chromosome. The CDH status of each control sample is shown on the x axis. Relevant features of each CDH are highlighted on the y axis. A scaled tag index was calculated for each 1 Mb bin; here the CDH haplotypes are scaled to their relative lengths. **B**. Stepwise manner of differentiating Ab10 from K10L2 (left) and general workflow for identifying CDHs using scaled tag index bins (right). Check indicates present, x indicates absence.

**Figure 2.**
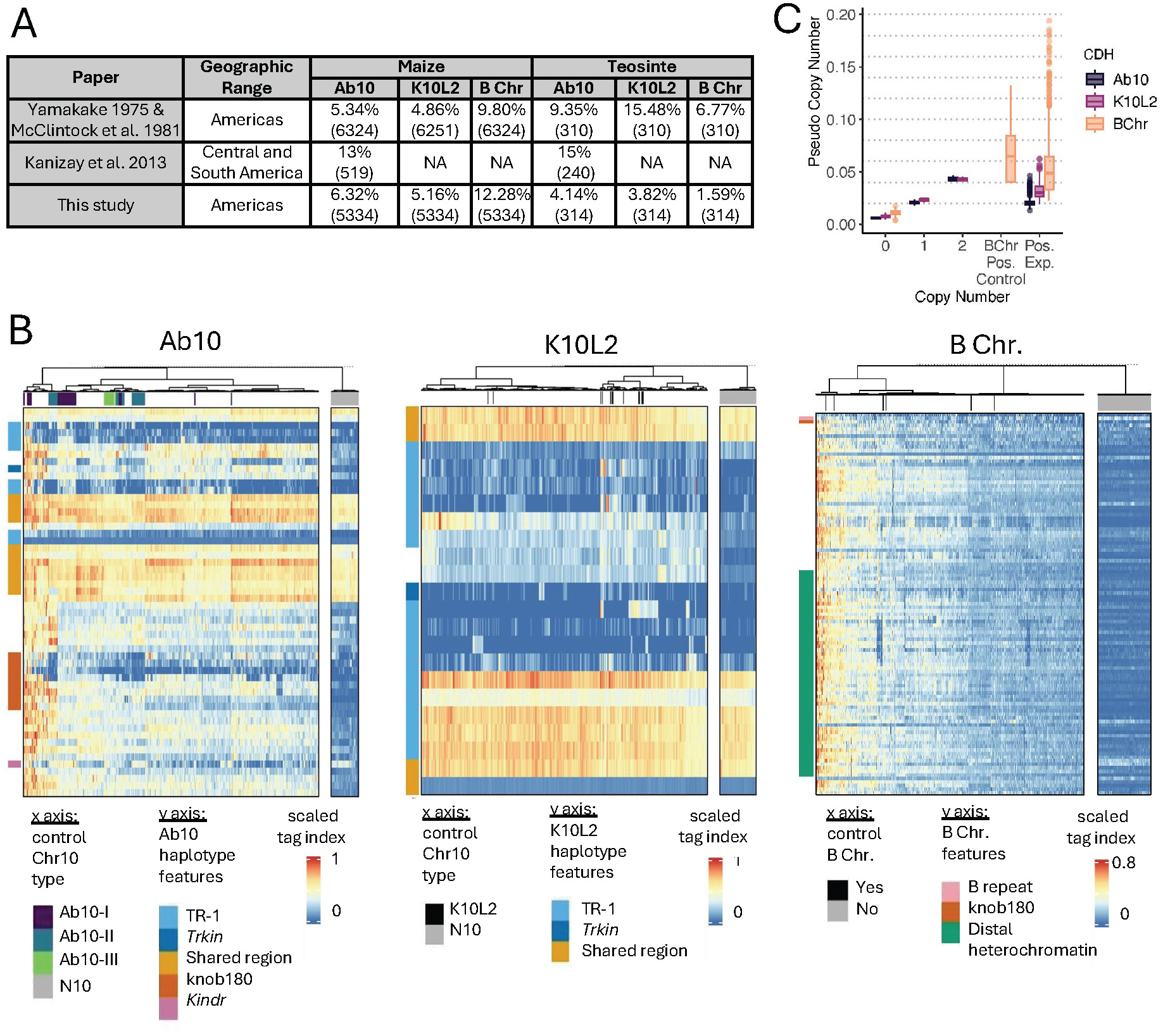
Detection of CDHs in experimental samples. **A**. Summary table for previous studies of CDH distribution as well as the results from the work presented here for comparison. Numbers in parentheses indicate the number of individuals surveyed. **B**. Scaled tag index for all CDH positive lines and N10 controls. CDH-positive controls and CDH-positive experimental samples are intermixed: those from controls are indicated by colors in the bars below the dendrograms (that show relatedness), those with no color are experimental samples. The N10 control samples are plotted separately. **C**. Pseudo copy number of all 3 CDHs in control and experimental samples. B Chr Pos. Control refers to samples that are B chromosome positive via PCR (copy number was unknown). Pos. Exp. refers to all CDH positive experimental samples identified in public GBS data. Dotted lines indicate approximate values for one copy as determined by the relationship between 1 and 2 copies in control samples. These are shown as a guideline only and are not meant to imply true copy number.

We then went on to identify CDH in previously published GBS data from landraces, teosintes and inbreds [42–44] using a similar stepwise k-means clustering approach (Fig S1B). Our Ab10 detection pipeline cannot detect K10L2, and our K10L2 pipeline cannot distinguish Ab10 from K10L2. Therefore we ran the Ab10 pipeline first, and then ran samples called as negative through the K10L2 detection pipeline (Fig 1B). Similarly, to accurately differentiate high copy B chromosome lines from low copy lines, we ran two different pipelines in sequence (see Methods). To ensure that clustering was driven by our control samples, we used roughly equal numbers (CDH positive and CDH negative) and randomly introduced experimental samples so that they were no more than 25% of the control samples (see Methods). Every sample was assayed for each CDH 125x to gain an estimate of call confidence. We obtained confident calls (>95% calls) for more than 99% of samples assayed for each CDH (Fig 2B, Fig S1B).

When developing our set of Ab10 control samples we had included isolates from three major cytological types, known as Ab10-I, Ab10-II, and Ab10-III, that differ in the appearance of the major knobs on the haplotype [45]. We developed a random forest model to detect Ab10 type from this control data. When this model was applied to samples with unknown Ab10 types, it became clear that there is more diversity in natural Ab10 samples than is present in our three control types (Fig 1A, Fig 2B). Using a confidence threshold that maintained the visual differences between types (see Methods), only 47 of 352 experimental Ab10-positive samples were classable (Fig S2A). To explore the variation in Ab10 type further, we performed a PCA on the scaled tag index of diagnostic regions. Plotting the first two principal components, accounting for 50% of the variation, revealed a nearly uniform distribution of the samples suggesting extensive genetic variation among Ab10 types (Fig S2B). Ab10-I and Ab10-II are known to recombine with each other in experimental populations [40], and some of the tag index patterns suggest recombinants are also present in natural populations (Fig S2A).

### Frequencies of three CDHs in maize

Our data on the frequencies of maize CDH in landraces are similar to what was previously reported by Kato, McClintock and Blumenschein based on their analysis of meiotic pachytene chromosomes ([19,20], Fig 2A). We observed Ab10 at a frequency of 6.32%, K10L2 at 5.16%, and the B chromosome at 12.28% on a per-plant basis. Each CDH occurs across the entire range of the landrace accessions assayed (Fig 3). We observed substantially fewer CDH in teosintes than was previously reported [19,20], which likely reflects the fact that the teosinte lines we assayed came from a relatively small number of accessions (Table S1). Ab10 and the B chromosome as well as K10L2 and the B chromosome occur together roughly as frequently as expected by chance (Table S1).

**Figure 3.**
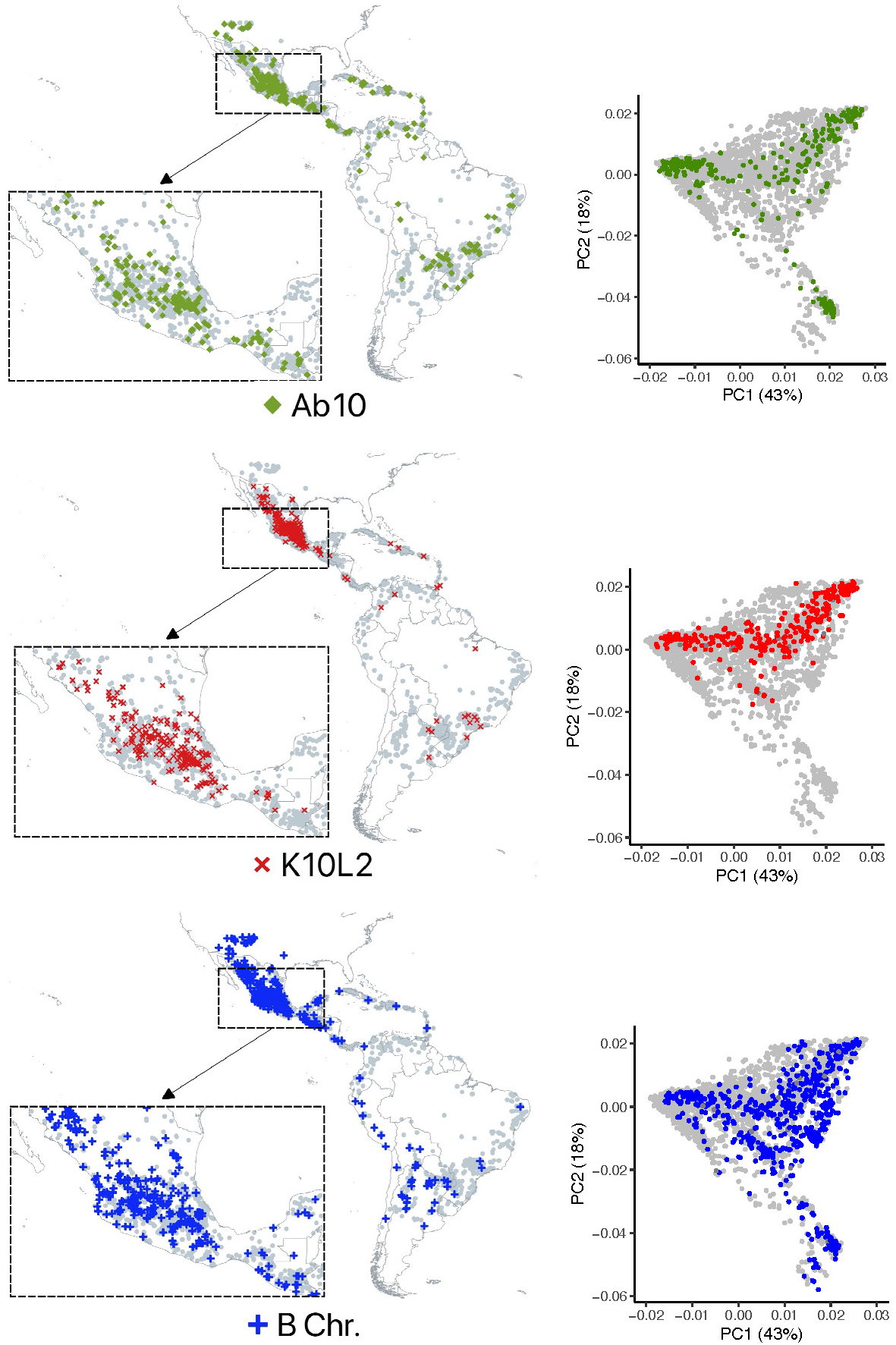
Maps and PCA plots of maize landraces assayed as CDH negative or positive. For each CDH, a location map is shown (left) and a plot of the first 2 principal components of population structure based on whole genome SNP profiling (right). Each grey dot indicates the location of a landrace that was assayed. Those with a CDH are highlighted in colors. The numbers in parentheses indicate the percent of variation that principal coordinates account for.

We also identified all three CDHs in inbred maize lines [44], though at low frequencies (Table S1, Table S2). There were 2 inbreds that scored positive for Ab10, 23 that scored positive for K10L2 and 16 that scored positive for the B chromosome (Table S2). K10L2 had previously been detected in an inbred line [23], but Ab10 and the B chromosomes were thought to be absent from inbred lines [18,28]. To confirm that the GBS genotyping was correct, we obtained seeds from six of the inbred lines – two that scored positive for each CDH. The seeds were from the Germplasm Research Information Network which maintains bulked samples derived from multiple ears. This is done to maintain any residual genetic diversity. We found that all six of the inbred lines contained individuals that scored positive for the CDH by PCR as well as individuals that scored negative by PCR (Table S2). While these lines are presumed to be pure breeding for most of the genome, the CDH chromosomes are still segregating for presence or absence. The fact that very few inbreds carry CDH, and many of those that do are incompletely inbred, supports the view that all three CDHs have deleterious fitness consequences.

### Relationship of three CDH in maize to genetic variants

We went on to examine the relationship between each CDH and genetic variation in the genome. Using the GBS data from maize landraces, we identified high confidence SNPs that did not overlap any CDH or transposable element, resulting in ∼50,000 usable SNPs (Fig S3). These data allowed us to identify population structure within our accessions (Fig 3) and perform a genome wide association study (GWAS) on high confidence SNPs that associate with each CDH. Given our binary traits and relatively large number of individuals we chose to impose a stringent significance threshold of 5×10-8.

We found that Ab10 was positively associated with seven SNPs on chromosomes 3, 4, 8, and 9 and negatively associated with four SNPs on chromosome 3, and 10. A highly associated SNP on chromosome 9 is likely an alignment artifact, as it occurs within a sequence that has homology to the Ab10-I v1 reference [41] and the p value is similar to what we observed for SNPs that are tightly linked to a CDH (Fig S4); it was not used in subsequent analysis. It is also possible that two other SNPs that are positively associated with very low p values (Ab10 Chr8 SNP1 and B chromosome Chr3 SNP1) are artifacts of alignment from CDH reads that differ significantly from our references; however in the absence of other evidence we have left them in the analysis. K10L2 was positively associated with five SNPs and negatively associated with two SNPs (Fig 4A). The B chromosome was positively associated with two SNPs and B chromosome copy number was not associated with any SNPs (Fig 4A). Of the total 20 associated SNPs, 13 overlap annotated genes (Table S3). Henceforth we refer to loci associated with each CDH as putative genetic modifiers for ease of interpretation.

**Figure 4.**
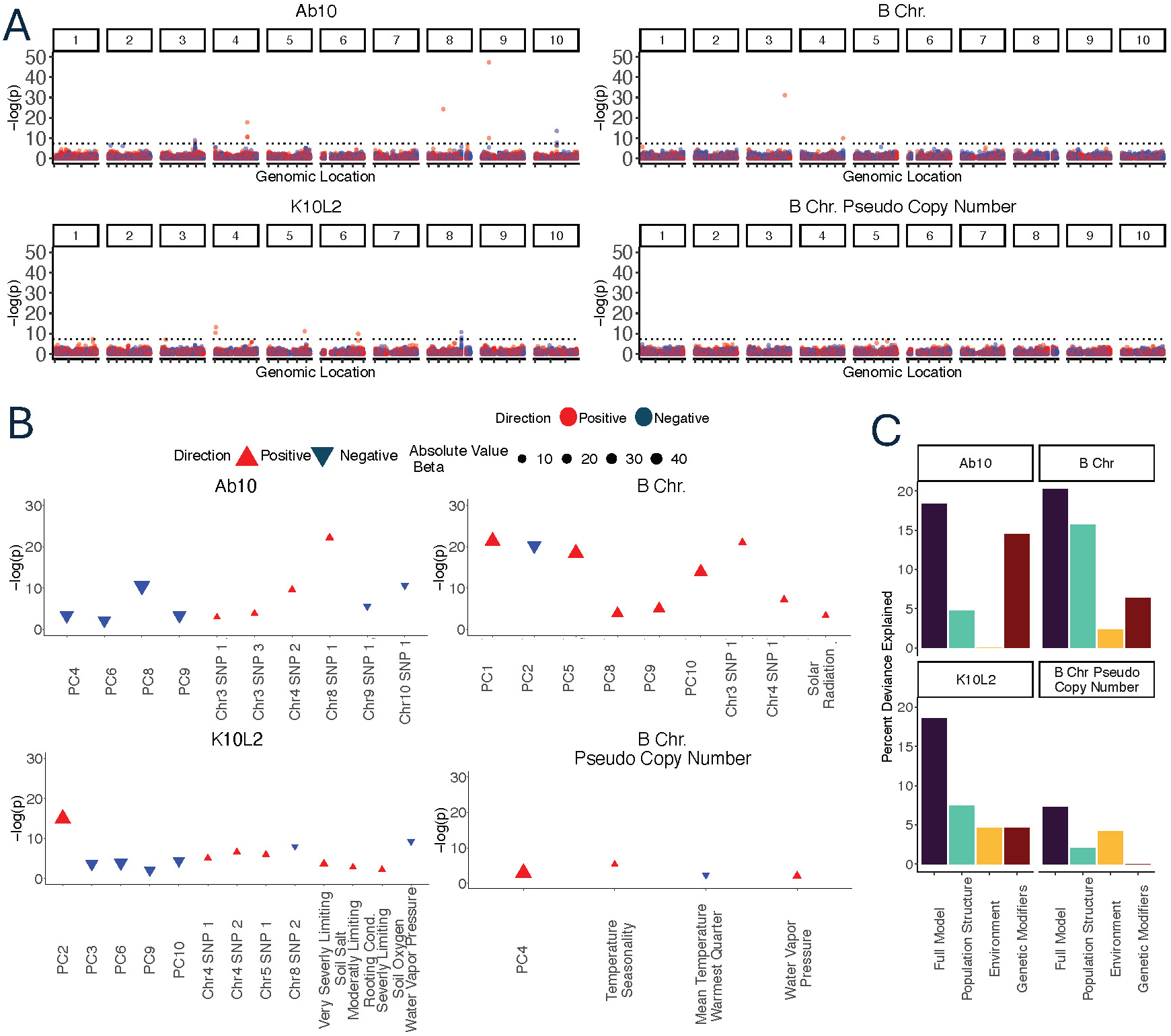
Relationship of CDH to the environment and genetic loci. **A**. Manhattan plots of SNPs that associate with each CDH and B chromosome copy number. Dotted black line indicates a p value of 5×10^-8^. The Ab10-associated SNP with very high p value (-(log)p close to 50) on chromosome 9 may be an artifact of read alignment to the Ab10 haplotype itself. **B**. Plots of fully simplified generalized linear models for each CDH including population structure, genetic loci, and environmental variables. Color and orientation of the triangle indicates whether a SNP is positively or negatively associated with the CDH. The size of the triangle represents the effect size. Very Severely and Severely Limiting Soil Salt refers to growth limiting excess soil salts. Moderately Limiting Rooting Cond. refers to moderately growth limiting rooting conditions. The direction of the relationship between Ab10 and Chr3 SNP1 and Chr9 SNP 1 changes when considering each SNP in isolation (A) or with the other SNPs and environmental variables (B). **C**. Partitioned deviance of each model shown in B. The partitions do not sum to the full model due to shared variation among the partitions.

### Combined effects of genetic variants and environment on the distribution of CDHs

To test the impact of location and environment on CDH distribution, we chose maize landrace and teosinte lines that were confidently identified as CDH positive or negative and had GPS coordinates for their collection location [Ab10= 3718, K10L2=3421, B Chr= 3718, B Chr Copy Number=542] (Table S1). We first assessed the effects of elevation while also accounting for population structure. In contrast to previous studies [27,31,45], we found no relationship between elevation and Ab10 or the B chromosome (presence/absence or copy number, Fig S5). There was a weak positive relationship between K10L2 and elevation when considering population structure alone (Fig S5).

We went on to develop models to test the effects of specific climatic variables [46] and soil conditions [47]. We began each model using the top 10 principal coordinates of population structure, as well as elevation, mean temperature of warmest quarter, precipitation of warmest quarter, temperature seasonality, precipitation seasonality, average annual solar radiation, average annual wind, average annual water vapor pressure, soil nutrient availability, soil rooting conditions, soil oxygen availability to roots, soil excess salts, and soil toxicity. For the B chromosome model, we also included Ab10 and K10L2 presence/absence to test whether they are independently distributed. For each CDH we generated a simplified model by removing variables that were not significantly associated (p value greater than 0.01) one at a time until all variables were significantly associated (p value less than 0.01). The presence or absence of Ab10 or K10L2 did not significantly affect the distribution of the B chromosome, consistent with expectations based on overlapping percentages (Table S1). There was no significant association between elevation and Ab10, K10L2, or the B chromosome when considering both population structure and climatic variables (Fig S6).

We then added genetic modifiers to the model, so as to include population structure, genetic modifiers and the environment (Fig 4). We calculated the amount of deviance explained by the full model and each of the variable classes [population structure, environment, and genetic modifiers] separately (Fig 4C). After accounting for all variables, only 12 of the original 20 SNPs showed significant associations. Due to interactions among the variables (which we did not pursue here), the deviance explained by each class of variables individually does not sum to the deviance explained by the full model.

#### Ab10

Ab10 significantly associates with six genetic modifiers and four principal components of population structure accounting for 18.4% of the deviance (Fig 4B,C). The environment seems to have no effect on the distribution of Ab10.

#### K10L2

K10L2 significantly associates with five principal components of population structure, four genetic modifiers, and four environmental factors accounting for 18.5% of the deviance (Fig 4B,C). It is associated with lower water vapor pressure. Further we found that K10L2 was overrepresented in poor quality soil specifically with respect to excess soil salts, soil rooting conditions, and soil oxygen (Fig 4B).

#### B Chromosome

The B chromosome significantly associates with six principle components of population structure, two genetic modifiers and one environmental variable accounting for 20.2% of the deviance. The B chromosome is more likely to occur in regions with higher solar radiation (Fig 4B,C).

B chromosome copy number is associated with one principle component of population structure, and three environmental variables accounting for 7.25% of the deviance. Specifically, B chromosome copy number is higher in environments with higher temperature seasonality and water vapor pressure but a lower mean summer temperature.

## DISCUSSION

In this work we used GBS data to identify chromosomal drive haplotypes in over 10,000 maize and teosinte accessions with the aim of better understanding how drive haplotypes interact with each other and the environment. While GBS was originally developed as a method to score SNPs [48], our approach using k-means clustering illustrates that the low coverage sequence data can also be used to identify large haplotypes such as CDHs. The method can accurately identify Ab10, K10L2 and B chromosomes, as well as variants that differ from the reference haplotypes that may be worthy of further study (Fig S2).

Although all three CDH can be transmitted at super-Mendelian levels, they are only found in natural populations at low frequencies, indicating that their distributions are limited by genetic or environmental factors. There is little doubt that the fitness consequences of the drivers themselves are major limiting factors. Lines carrying Ab10 have reduced seed number and weight, and the B chromosome causes sterility when present at high copy numbers [17,28]. However, at least in the case of Ab10, modeling shows that the known fitness defects are not sufficient to explain the low observed frequencies of Ab10 [21]. Here we assessed the importance of population structure, genetic modifiers, and the environment to the distribution of Ab10, K10L2, and B chromosomes, as well as their interactions with each other.

The environment appears to have little effect on the distribution of any of the maize chromosomal drivers. Prior data suggested that the B chromosomes and Ab10 might occupy different altitudinal clines [27,31,45]. However, after controlling for population structure, we found no correlation with elevation for either driver, and that Ab10 and the B chromosome occur together as frequently as expected by chance (Table S1). There has been an assumption in the prior literature that since both the B chromosome and Ab10 increase genome size, they should be selected against at higher altitudes where smaller-genome lines are more fit [33]. While this may be true, the level of selection may be weaker than is commonly assumed. Recent results suggest that in the large-genome maize plant, a gain of 14 Mb results in a 0.1% reduction in yield [49]. By this reasoning, a single ∼30 Mb knob [13], the ∼85 Mb Ab10 haplotype [14], or the ∼150 Mb B chromosome would be expected to result in an ∼<1% drop in yield, which may not be sufficient to counteract the selfish properties of these powerful drivers.

We used a GWAS approach to identify potential genetic modifiers that impact the distribution of the maize CDH. At the outset, we anticipated that most of the modifiers would be negatively associated, and represent potential suppressors that reduce the efficiency of the drivers. Extensive prior literature suggests that such suppressors are likely to evolve when the fitness burden is high [2,29,36,37]. For instance, we anticipated identifying SNPs that are linked to alleles that suppress the preferential fertilization of the egg by B chromosome-carrying sperm [8]. However, only 3 of the 12 associated loci showed such a negative relationship (Fig 3B, significant SNPs after accounting for environment). The remaining 9 SNPs were positively associated. While enhancers of meiotic drive are known to exist, they are not expected to evolve in positions that are unlinked to a driver [50]. This surprising outcome suggests that the maize genome may be adapting to the presence of chromosomal drivers with unlinked alleles that reduce their negative fitness consequences. This would be similar, for example, to genetic modifiers that reduce disease severity in humans [51], or mutations that bypass the phenotype of otherwise lethal mutations in yeast [52].

Taken together, our data suggest that the major limit to the spread of Ab10, K10L2 and the B chromosome are the fitness defects associated with the drivers themselves at high copy numbers. The fact that modeling based on the known fitness defects associated with Ab10 (reduced seed number, reduced seed weight and mildly reduced pollen viability [17]) does not predict the observed low frequencies suggests that there are additional fitness defects that have not yet been accounted for [21]. One way to better assess the fitness defects would be to start with the few (near)-inbred lines that we show are still segregating for the major chromosomal drivers (Table S2). Further self crossing should make it possible to identify sibling inbreds that either do or do not carry the CDH, and measure a broader array of fitness and fertility variables, including, for instance, seed germination and survival from seedling to reproductive stages [21].

## METHODS

### GBS Sequencing Controls

Our control GBS data were obtained from two different sources, Cornell and CD Genomics. For the Cornell dataset, plants known to be heterozygous for Ab10-I-MMR or Ab10-II-MMR were self crossed to create populations segregating for either Ab10 structural variant. Ab10 was marked by an allele of the *colored1* gene (*R1*) which makes the kernels purple (Table S4). There is an approximately 2% chance of recombination between Ab10 and *R1* [53]. We extracted genomic DNA from plants grown from purple seeds (likely Ab10 positive) and colorless seeds (likely N10 homozygotes) using a CTAB extraction protocol [54] (Ab10-I- MMR=41, Ab10-II-MMR= 37, N10=16). Using these DNA samples, GBS libraries were prepared and sequenced on an Illumina HiSeq 2000 in accordance with [48] by the Genomic Diversity Facility, Cornell University (this facility is no longer in operation).

For the CD genomics dataset, we grew plants from 49 Ab10 controls from 11 genetic backgrounds, 13 K10L2 controls from 2 genetic backgrounds, 18 B chromosome controls from 5 genetic backgrounds, and 18 no CDH controls from 7 genetic backgrounds (Table S4). We first verified that the controls were CDH positive or negative by extracting DNA using CTAB extraction [54] and performing PCR for *Kindr*, *Trkin*, or the B repeat (Table S5). We then sent leaf tissue to CD Genomics (Shirley, NY) who extracted DNA using QIAgen DNeasy Plant Kits. They prepared GBS libraries as described in [48] with minimal modification. Basically this involved digesting DNA with ApeKI (New England Biolabs, Ipswich, MA), adding barcoded adapters, and sequencing the libraries on an Illumina NovaSeq6000 using a 150×2 paired-end sequencing protocol.

After receiving the data we identified several lines that appeared to be misclassified based on scaled tag index k-means clustering (W23_AB10-I.11.DC1, W23_AB10-I.13.DC1, W23_AB10-II.36.DC1, W23_N10.14.DC1, NSL-2833_B-Chrom.2.DC2, B542C_L289_B- Chrom.1.DC2). W23_AB10-I.11.DC1, W23_AB10-I.13.DC1, W23_AB10-II.36.DC1, and W23_N10.14.DC1 were likely recombinants, but this could not be verified by PCR as we no longer had the samples and were excluded from further analysis. NSL-2833_B-Chrom.2.DC2 and B542C_L289_B-Chrom.1.DC2 were re-genotyped and reclassified as having no B chromosomes.

### Obtaining GBS Data

We obtained GBS sequence reads from the authors of three prior publications [42–44]. These data were generated following the protocol of [48]. The data from [43] were in the format of demultiplexed qualified reads; we converted them to a format usable for TASSEL using custom R v4.3.1 code and barcode faker [55]. The data from each plant described in [43] was split into approximately 4 libraries as technical replicates, and these were summed during analysis (see below).

### K-means Clustering of Controls

We first established that it was possible to differentiate Ab10, K10L2, and N10 from each other as well as B chromosome presence/absence using GBS data. We began by mapping the full set of control GBS data to the B73-Ab10 v2 [14] genome with the B chromosome appended [24] and the CI66 inbred genome carrying the K10L2 haplotype [14] using TASSEL v5.2.44 [55] and BWA v0.7.17 [56]. Using TASSEL v5.2.44 [55], we obtained the coordinates of each tag and the number of associated reads in each sample for both the B73-Ab10/B-Chromosome assembly and the CI66-K10L2 assembly. We converted the alignments to a bed file using samtools v0.1.20 [57], and bedtools v2.29.2 [58]. For each assembly, we summed the tag counts for all technical replicates per biological individual for [43]. Unless otherwise noted all further steps were carried out using custom R v4.3.1 code. In order to normalize across libraries of varying size, we calculated reads per million for each tag in each individual sample. We calculated the minimum proportion of missing data for blank samples (where no genomic DNA was added; this represents sequencing background), and subtracted 0.001. We then removed any sample with more missing data than this cut off, as well as any tag with a BWA mapping quality of less than 20. We verified that all datasets were affected similarly by these filters and extracted all tags on each CDH. We then calculated the tag index in non overlapping 1 Mb bins across all CDHs (sqrt(c) + d), where c is the count of tags mapped to that bin and d is the sum of the read depth of all tags in that bin (Fig S1A). Then we visualized control samples with known CDH status in a min/max scaled heat map of the tag index. We found that the CDH positive and negative lines were visually very distinct (Fig 1A). We did not observe any visual distinction between our two sets of control data (the Cornell and CD genomics datasets), indicating that this method is robust to differences in sequencing. This is important as the experimental data set is pooled from multiple data sources.

We then established that we could correctly and automatically detect CDH presence or absence in our control data set. We chose to use an iterative k-means clustering method on the scaled tag index. The entire pipeline outlined below was performed using custom R v4.3.1 code unless otherwise stated. We selected only the CDH-specific portions, which were the regions that showed stark differences between CDH positive and negative controls (Fig 1A). For each CDH we had high and low copy number controls. For Ab10 and K10L2, the high copy number controls were homozygous plants with two copies of the CDH, and low copy number controls were heterozygous plants with one copy of the CDH. For the B chromosome, the copy number was unknown and they were divided into high and low copy number controls by visual comparison of the min/max scaled tag index heat maps (Fig 1A). We analyzed the high and low copy number controls separately in order to ensure that clustering was based on the distinction between presence and absence rather than copy number. First, we split our group of control samples into three randomly selected groups. On each subsample we performed k-means clustering (k=2). If a cluster was composed of at least 80% CDH (Ab10, K10L2, B chromosome) or non CDH (N10, no B chromosome) samples it was assigned as such (this is the naming step in Fig 1B). The k-means cluster assignment was then compared to the true CDH status of that sample in order to determine if the k-means clustering assigned the sample correctly. This was repeated 100 times for each sample, where each iteration involved a different, randomly selected set of control individuals (Fig 1B). For the B chromosome low copy number model we didn’t have adequate samples to break them into three subsamples so they were clustered as a single group. Using this method we were able to correctly identify the CDH status of all of the control samples 100% of the time, regardless of where the GBS data were acquired (either from Cornell or CD genomics).

### Use of K-means Clustering on Experimental Samples

Having established that the method correctly identifies each CDH in control data, we then extended it to our experimental samples [42–44]. We identified the chromosome 10 CDHs and the B chromosomes in two separate workflows before finally estimating their copy number. The entire pipeline outlined below was performed using custom R v4.3.1 code unless otherwise stated.

The general approach was to select roughly equal numbers of the appropriate controls (positive and negative) for each CDH and then randomly add a small number of experimental samples. For Ab10 and K10L2, the number of experimental samples added was 25% of the number of controls, for the B chromosome, the number of experimental samples added was 10% of the number of controls. Then we performed k-means clustering (Fig S1B). If a cluster was composed of at least 80% CDH (Ab10, K10L2, B chromosome) or non CDH (N10, no B chromosome) samples it was assigned as such. We verified that all control samples were correctly identified. If they were not, we repeated the k-means clustering until all controls were correctly identified. We then assigned all experimental samples to the class of their k-means cluster. We repeated this workflow until all experimental samples had undergone one round of k-means clustering. Then we repeated the entire process 125 times to obtain 125 independent calls for the CDH class per experimental sample. To make the final CDH class calls, we required that the experimental sample be called the same class 95% of the time. All other samples were labeled ambiguous.

Our Ab10 pipeline cannot distinguish K10L2 from N10, while our K10L2 pipeline cannot distinguish Ab10 from K10L2. Therefore we employed them one after the other. We ran the Ab10 pipeline first and identified 394 Ab10 positive samples. We then isolated the samples called as N10 and ran the K10L2 pipeline, identifying 310 K10L2 positive samples. We plotted all the Ab10 and K10L2 positive samples in single heat maps with ward.D clustering (Fig 2B).

The variability in B chromosome copy number in experimental samples sometimes caused our k-means clustering pipeline to fail (lines with many copies of the B chromosome sometimes formed their own cluster). Therefore we used a two-step process. First we extracted all high copy number experimental samples using high copy number controls. Then we took all samples not identified as B chromosome positive in the high copy number iteration and ran them through the same pipeline using the low copy number B chromosome controls. In this way we were able to extract all B chromosome positive samples without introducing unnecessary variation in the k-means clustering. We plotted all B chromosome positive samples in a single heat map with ward.D clustering (Fig 2B).

### Random Forest Modeling

We then attempted to differentiate the Ab10 types within our experimental classes. We first trained a random forest model on 70% of the Ab10 control data with known types [17]. We checked the random forest model’s performance using the remaining 30% of the Ab10 control data. It correctly predicted type 100% of the time. We then applied the same random forest model to all of our experimental samples. We required that 65% of decision trees call the same Ab10 type; if less than 65% called the same type, the haplotype was classified as ambiguous (Fig S2A). We selected this confidence threshold as it preserved the visually apparent differences between types when plotted as a heat map (Fig 1A). However, only 11.9% of Ab10 samples were classable in this manner. To better explore Ab10 types we extracted the bins with the highest mean decreasing Gini in the random forest model, meaning the model suffered the most when these variables were excluded, and performed a principal coordinate analysis (Fig S2B).

### Estimating the Copy Number of CDHs

To estimate the copy number for each CDH we needed an estimate of what the tag index of a single copy gene was. To this end we lifted over annotations from the B73 v5 reference [12] onto both the Ab10 v2 and CI66 K10L2 assemblies [14] using liftoff v1.6.3 [59]. We then extracted all single copy core genes, and calculated their tag index in 1 Mb bins (Fig S1A). The 1 Mb bins were composed of 1 Mb of single copy core gene sequence and not genomic coordinates. Then we calculated the average tag index across all single copy core gene bins for each sample. We divided the average CDH specific tag index value by that sample’s average single copy core gene tag index value. We refer to the ratio of CDH/single copy core gene tag index as pseudo copy number (Fig 2C).

### PCR verification of CDHs in inbred lines

The frequency of CDHs varies in open pollinated populations like landraces and teosinte and are rarely if ever fixed [23,25,45]. Thus while we may have scored one plant from a landrace as positive for a CDH, it is unlikely that the next plant we scored from the same population would have the CDH. However, we identified several maize inbred lines containing Ab10, K10L2, and B chromosomes (Table S1). We ordered two inbred lines called as positive for each CDH from the Germplasm Resources Information Network (GRIN), Ames IA. We extracted DNA using a CTAB extraction [54] and performed PCR for *Kindr*, *Trkin*, or the B repeat to verify the CDHs presence (Table S2, Table S5).

### GWAS

We generated artificial reference genomes with Mo17 [13] chromosome 1-10 and the Ab10 v2 haplotype [14], K10L2 haplotype [14], or the B chromosome [24]. For the Mo17 Ab10 and K10L2 reference genomes we used samtools v1.18 [57] to truncate Mo17 chr10 at the beginning of the *colored1* gene, which traditionally defines the beginning of the Ab10 and K10L2 haplotypes [18]. We then isolated the CDHs beginning at the *colored1* gene using samtools v1.18 [57] and SeqKit v0.16.1 [60] and appended them to the modified Mo17 genomes. For the B chromosome we left all Mo17 chromosomes intact and added the B chromosome [24]. We modified the key for all samples such that all technical replicates from [43] were read into a single biological sample. We then used TASSEL v5.2.44 [55] and BWA v0.7.17 [56] to align reads from all samples to the Mo17 + CDH references. We filtered mapped tags to a mapping quality of 20 using samtools v1.18 [57] and called SNPs using TASSEL v5.2.44 [55]. We then extracted SNPs on chromosomes 1-10 and the relevant CDH using bcftools v1.15.1 [57]. We isolated only maize landraces using bcftools v1.15.1[57] because we suspected genetic modifiers might be less frequent in inbred lines due to the low frequency of CDHs (Table S1).

Then we applied the following filters: Read depth >3 and <20, minor allele frequency >= 0.05, genotype quality >60, and a per sample missingness of 75% or less. We used BEAGLE v5.4 to impute missing data based on haplotypes found in our data [61]. We did not use a reference panel due to concerns about the maintenance of genetic modifiers for CDHs in inbred lines.

Using PLINK v1.9 [62] we removed plants with more than 10% missing data. We were left with ∼50,000 SNPs. We carried out a principal component analysis on the whole genome non-CDH SNPs to identify population structure in the data, and included the top 10 principal components in a genome wide association test as covariates using PLINK v1.9 [62]. For Ab10, K10L2, and B chromosome presence/absence we used a logistic regression and for B chromosome pseudo copy number we used a linear regression. We plotted the output using custom R v4.3.1 code.

GBS tags are just 64 bp [55]. We know that some regions of each CDH are homologous to chromosomes 1-10 [14,18,23]. It seemed possible that a GBS tag originating from a CDH could map to chromosomes 1-10 and create an erroneous association. We identified genes orthologous between the CDH and the Mo17 genome using OrthoFinder v2.5.5 [63] and removed any associated loci overlapping them using bedtools v2.31.0 [58]. Additionally we removed any SNP overlapping an annotated transposable element [13] using bedtools v2.31.0 [58]. Finally, we extracted 64 bp upstream and downstream of each SNP using Samtools v1.18 [57] and used BLAST v2.13.0 to compare the region to the CDH in the B73-Ab10 v2 [14], K10L2 [14], or B chromosome [24,64] references. We removed any SNP that had homology to a CDH of at least 62 bp with a percent identity of 80% or greater. The sequence surrounding Chr9 SNP 2 (29026173) had 83% percent identity to the Ab10 v1 reference [41], though there was no significant homology to the B73-Ab10 v2 reference [14]. We suspect this associated SNP is an artifact and excluded it from further analysis.

We repeated the above procedure using all the same criteria on SNPs on the CDH as a control for loci linked to the CDH (Fig S4). We removed SNPs that did not occur in at least 75% of the samples, so any locus specific to the CDH should have been removed. Loci present in the inverted shared region of Ab10 could exist in individuals carrying Ab10 or N10 allowing them to pass the missing data filters. While K10L2 also has a shared region it is not inverted [14] and is known to recombine with N10 [23] so while many SNPs passed the filtering we would expect fewer of them to strongly associate with K10L2 presence. The B chromosome does not have a shared region thus most SNPs on it occur in less than 75% of samples and were removed by the missing data filter.

### GLM models on All CDHs

We obtained climatic data from WorldClim2 [46] and soil quality data from the FOA harmonized world soil database [47]. We chose to begin each model with environmental variables known to be associated with either maize or the CDHs: elevation, mean temperature of warmest quarter, precipitation of warmest quarter, temperature seasonality, precipitation seasonality, average annual solar radiation, average annual wind, average annual water vapor pressure [46], soil nutrient availability, soil rooting conditions, soil oxygen availability to roots, soil excess salts, and soil toxicity [47,65]. For Ab10 we included B chromosome presence/absence (we could not include K10L2 because we cannot detect K10L2 in lines where Ab10 is present). For the B chromosome we included Ab10 and K10L2 presence/absence. We extracted environmental data for each collected sample using the raster package in R v4.3.1 [66]. For solar radiation, average annual wind, and average annual water pressure we used custom R v4.3.1 code to generate the average value for each location from monthly data [46].

We tested for collinearity between the environmental variables using custom R v4.3.1 code and found that elevation and soil nutrient availability had a greater than 70% correlation to other variables (Fig S7). We excluded soil nutrient availability from all starting models due to its collinearity. We chose to include elevation in the starting models because it has been investigated previously [32,45]. We also included the first 10 principal components of population structure generated as part of the GWAS. For the CDH presence/absence we used binomial family models. For B chromosome pseudo copy number we log transformed the variable to make it normally distributed and used a gaussian family model. From the largest model we performed stepwise model simplification using custom R v4.3.1 code. We ensured all models were robust to variable order. We tested model fit using the DHARMa R package before and after simplification [67]. We determined model fit was acceptable in all cases. Our models represented a relatively low proportion of the total deviance, thus we chose to apply an alpha value of 0.01.

To assess the relative contributions of population structure, genetic loci, and the environment we selected all significant loci from the GWAS using custom R v4.3.1 code. Because some loci had more than one alternate allele, we began by coding these as factors to check if any of the minor alleles had a significant relationship to any CDH. We found that they did not and removed them, coding the remaining minor alleles additively. We then performed stepwise model simplification. We plotted the results using custom R v4.3.1 code. To partition the variance from the entire model into population structure, genetic loci, and environmental variables we first used custom R v4.3.1 code to remove all missing data to ensure the null model was identical between all runs. Then we ran individual models with only the remaining population structure, genetic loci, or environmental variables. We calculated the amount of deviance explained by each of these models and plotted them using custom R v4.3.1 code.

### Data Availability

All code can be found at https://github.com/dawelab/CDH_Distribution_In_Maize. GBS data can be accessed at the SRA under <u>PRJNA1257855</u>.

## Acknowledgements

We thank Douda Bensasson for guidance on modeling, Brad Nelms for help developing the CDH detection method, and Robert Unckless and Jonathan Gent for helpful comments on the manuscript. We also thank Cinta Romay, Edward Buckler, and the International Maize and Wheat Improvement Center (CIMMYT) for providing the raw GBS reads. We are grateful to the Georgia Advanced Computing Resource Center for providing technical expertise. This work was supported by an NIH training grant (T32GM007103) and NSF fellowship to MJB (2236869) as well as an NSF grant to RKD (1925546).

---

**Table S1.** Summary table of all CDHs identified.

**Table S2.** Inbred lines that scored positive for CDHs, and PCR data from a subset.

**Table S3.** All CDH associated SNPs and Genes they overlap.

**Table S1.** Accessions and individuals assayed, and lines that show both B chromosome and Ab10 or B chromosome and K10L2.

**Table S5.** Primers used for genotyping.

**Figure S1.** Workflow diagrams. **A.** Workflow diagram for the generation of the tag index for CDHs and single copy core genes. **B.** Diagram of the workflow for detecting CDHs in experimental samples. Check indicates passing, x indicates failing.

**Figure S2.** Identification of Ab10 type. **A.** Min/Max scaled tag index for all Ab10 positive control and experimental samples. Ab10 types classified by the random forest model are plotted separately. Each group is ward.D clustered. The x axis shows individual samples, the y axis shows features of the Ab10 haplotype and the importance of each 1 Mb bin in determining Ab10 type in the random forest model (mean decreasing Gini). The RF confidence value indicates the proportion of decision trees that are called the predominant Ab10 type. **B**. A PCA of all the Ab10 positive samples scaled tag index with controls and their type indicated.

**Figure S3.** Location of SNPs used for GWAS. Numbers below the CDH name indicate the total number of SNPs.

**Figure S4.** Manhattan plots of SNPs that passed through filtering but lie within or close to a CDH. Numbers below the CDH names indicate the number of SNPs in the plot. Dotted grey line indicates a p value of 5×10^-8^.

**Figure S5.** Relationship of CDHs to elevation. N.S. indicates not significant.

**Figure S6.** Plots of simplified generalized linear models for each CDH including population structure and environmental variables (but not genetic modifiers). Shape color and orientation indicate the direction of the relationship to the CDH. Shape size represents the effect size.

**Figure S7.** Correlation matrix for selected environmental variables.

